# LncRNA Mrhl orchestrates differentiation programs in mouse embryonic stem cells through chromatin mediated regulation

**DOI:** 10.1101/713404

**Authors:** Debosree Pal, C V Neha, Utsa Bhaduri, Zenia, Subbulakshmi Chidambaram, M.R.S Rao

## Abstract

Long non-coding RNAs (lncRNAs) have been well-established to act as regulators and mediators of development and cell fate specification programs. LncRNA Mrhl (meiotic recombination hotspot locus) has been shown to act in a negative feedback loop with WNT signaling to regulate male germ cell meiotic commitment. In our current study, we have addressed the role of Mrhl in development and differentiation using mouse embryonic stem cells (mESCs) as our model system of study. We found Mrhl to be a nuclear-localized, chromatin-bound lncRNA with moderately stable expression in mESCs. Transcriptome analyses and loss-of-function phenotype studies revealed dysregulation of developmental processes and lineage-specific genes along with aberrance in specification of early lineages during differentiation of mESCs. Genome-wide chromatin occupancy studies suggest regulation of chromatin architecture at key target loci through triplex formation. Our studies thus reveal a role for lncRNA Mrhl in regulating differentiation programs in mESCs in the context of appropriate cues through chromatin-mediated responses.

## Introduction

Since their discovery, embryonic stem cells (ESCs) and their potential in regenerative medicine have been widely established. They have been programmed to form cone cells for the treatment of age-related macular degeneration (Zhou et al., 2015), used for the treatment of spinal cord injury patients (Shroff & Gupta, 2015), differentiated into cardiomyocytes for cardiac tissue injury repair (Shiba et al., 2012) or into insulin secreting β-cells for treating Type II diabetes (Bruin et al., 2015; Salguero-Aranda et al., 2016). Their self-renewal characteristics coupled with their capacity to differentiate into cell types of origin pertaining to all the three germ layers make them an attractive model system to study queries related to development (Evans & Kaufman, 1981; Martin, 1981). This entails that the molecular pathways and players contributing to the physiology of stem cells be thoroughly characterized to explore their therapeutic potential to the fullest.

Long non-coding RNAs (lncRNAs) have been classified as non-coding RNAs that are >200nt in length and have been well-established to regulate diverse physiological phenomena from development to disease. Although they are similar to mRNAs in aspects such as transcription by RNA Pol II, 5’ capping, 3’ polyadenylation, splicing and possession of histone modification signatures of active transcription across their promoters and gene bodies (Guttman et al., 2009), they differ in other aspects such as low abundance, high tissue-specificity, lower stability and lesser evolutionary conservation (Ard, Allshire, & Marquardt, 2017; Mercer, Dinger, & Mattick, 2009). Furthermore, they can be localized either in the nucleus or in the cytoplasm to perform their diverse regulatory functions such as regulation of chromatin architecture (Rinn et al., 2007; Zhao, Sun, Erwin, Song, & Lee, 2008), genomic imprinting (Pandey et al., 2008; Ripoche, Kress, Poirier, & Dandolo, 1997; Sleutels, Zwart, & Barlow, 2002), competitive endogenous RNAs (Cesana et al., 2011; Lu et al., 2016), natural antisense transcripts (Faghihi et al., 2008; Ohhata, Hoki, Sasaki, & Sado, 2008), Staufen-mediated mRNA decay or modulation of protein activity (Arun, Akhade, Donakonda, & Rao, 2012; N. Lin et al., 2014; Marchese et al., 2016). LncRNAs are being increasingly understood in terms of regulation of the nuclear architecture through the formation of speckles and paraspeckles and tethering of distal chromosomes (Caudron-Herger et al., 2015; Caudron-Herger & Rippe, 2012; Hacisuleyman et al., 2014; Jacob, Audas, Uniacke, Trinkle-Mulcahy, & Lee, 2013; Shevtsov & Dundr, 2011; Souquere, Beauclair, Harper, Fox, & Pierron, 2010).

In embryonic stem cells (ESCs), genome-wide studies and functional analysis of individual lncRNAs have shown them to participate in maintaining pluripotency (Bergmann et al., 2015; Guttman et al., 2011; Sheik Mohamed, Gaughwin, Lim, Robson, & Lipovich, 2010; Z. Sun et al., 2018; Tu, Tian, Cheung, Wei, & Lee, 2018) as well as in orchestrating differentiation and cell fate specification programs (Flynn & Chang, 2014; Klattenhoff et al., 2013; Ulitsky, Shkumatava, Jan, Sive, & Bartel, 2011), in association with transcription factors, chromatin modifiers and RNA binding proteins. Interestingly, a few of these lncRNAs perform multiple roles as a function of the cellular contexts and interaction partners. LncRNA Gomafu/Miat/Rncr2 is involved in maintaining pluripotency of mouse ESCs (mESCs) (Sheik Mohamed et al., 2010), specification of the oligodendrocyte lineage in neural stem cells (Mercer et al., 2010) and osteogenic lineage differentiation in adipose-derived stem cells (Jin et al., 2017). LncRNA Tuna has been implicated in maintaining pluripotency of mESCs as well as their differentiation into the neural lineage through forming a complex with PTBP1, hnRNP-K and NCL that occupies promoters of *Nanog*, *Sox2* and *Fgf4* (N. Lin et al., 2014). LncRNA Tsx has also been shown to be involved in the maintenance of mESCs, pachytene spermatocytes in testes and regulation of cognition and behaviour in mice (Anguera et al., 2011). These examples suggest the diversity and the context-dependant regulatory functions of lncRNAs in stem cell physiology and demand further investigations on the roles of lncRNAs in ESCs.

LncRNA Mrhl (meiotic recombination hotspot locus) was discovered in our laboratory and has been extensively studied in the context of male germ cell meiotic commitment. It is a 2.4 kb long, sense, intronic and single-exonic lncRNA, encoded within the 15th intron of the *Phkb* gene in mouse (Nishant, Ravishankar, & Rao, 2004) and is syntenically conserved in humans (Fatima, Choudhury, Divya, Bhaduri, & Rao, 2018). It has been shown to act in a negative feedback loop with WNT signaling in association with its interaction partner p68 to regulate meiotic progression of type B spermatogonial cells (Akhade, Dighe, Kataruka, & Rao, 2016; Arun et al., 2012). Genome-wide chromatin occupancy studies have revealed regulation of key genes involved in spermatogenesis and WNT signaling by Mrhl in a p68 dependant manner, one of them being the transcription factor SOX8 (Akhade, Arun, Donakonda, & Rao, 2014). Subsequent studies have delineated the mechanisms of WNT mediated down regulation of Mrhl through the recruitment of the co-repressor CTBP1 at the promoter of Mrhl and regulation of SOX8 by Mrhl through MYC-MAD-MAX complexes (Kataruka, Akhade, Kayyar, & Rao, 2017), suggesting an intricate network of Mrhl and associated proteins acting to orchestrate the process of male germ cell meiosis.

In purview of the aforesaid, we have addressed the role of lncRNA Mrhl in mESCs to understand it as a molecular player in development and differentiation. We demonstrate through transcriptome studies that depletion of Mrhl in mESCs leads to dysregulation of 1143 genes with major perturbation of developmental processes and genes including lineage-specific transcription factors (TFs) and cell adhesion and receptor activity related genes. mESCs with stable knockdown of Mrhl displayed aberrance in specification of ectoderm and mesoderm lineages with no changes in the pluripotency status of the cells, consistent with our transcriptome data. Genome-wide chromatin occupancy studies showed Mrhl to be associated with ~22,000 loci. To decipher chromatin-mediated target gene regulation, we overlapped the two datasets and found key developmental TFs such as RUNX2, POU3F2 and FOXP2 to be directly regulated by Mrhl possibly through RNA-DNA-DNA triplex formation. Our study delineates lncRNA Mrhl as a chromatin regulator of cellular differentiation and development genes in mESCs, probably acting to maintain the cells in a more primed state readily responsive to appropriate differentiation cues.

## Results

### Mrhl is a nuclear-localized, chromatin bound moderately stable lncRNA in mESCs

We analyzed poly (A) RNA-Seq datasets from the ENCODE database and observed that Mrhl is expressed predominantly in the embryonic stages of tissues of various lineages (**Figure 1-supplementary figure 1**). In the brain, heart and lung, Mrhl is expressed all throughout different stages of embryos whereas its expression is almost nil in the postnatal stages. However, in the liver and the kidney, Mrhl expression is down regulated but not completely abrogated in the postnatal stages. From E8.5 onwards, the mouse embryo undergoes a surge of differentiation, cell specification and organogenesis phenomena. Our data analysis suggested that Mrhl might have a selective role to play in these processes in the context of mouse embryonic development. To address this, we used mESCs as our model system of study. RNA FISH revealed Mrhl to be expressed primarily in the nulcei of mESCs (Figure 1A). Biochemical fractionation further validated Mrhl to be present in the nuclear fraction, specifically the chromatin fraction in mESCs (Figure 1B, C). We next addressed if Mrhl was actually associated with the chromatin for which we performed H3 ChIP followed by qPCR and observed significant enrichment of Mrhl in H3 bound chromatin (Figure 1D). To further understand the functional relevance of Mrhl in mESCs, we performed an assay for RNA half-life and we found Mrhl to display moderate stability with a half-life of 2.73 hours (Figure 1E). Our observations herewith prompted us to investigate further the functional roles of Mrhl in mESCs.

**Figure 1:**
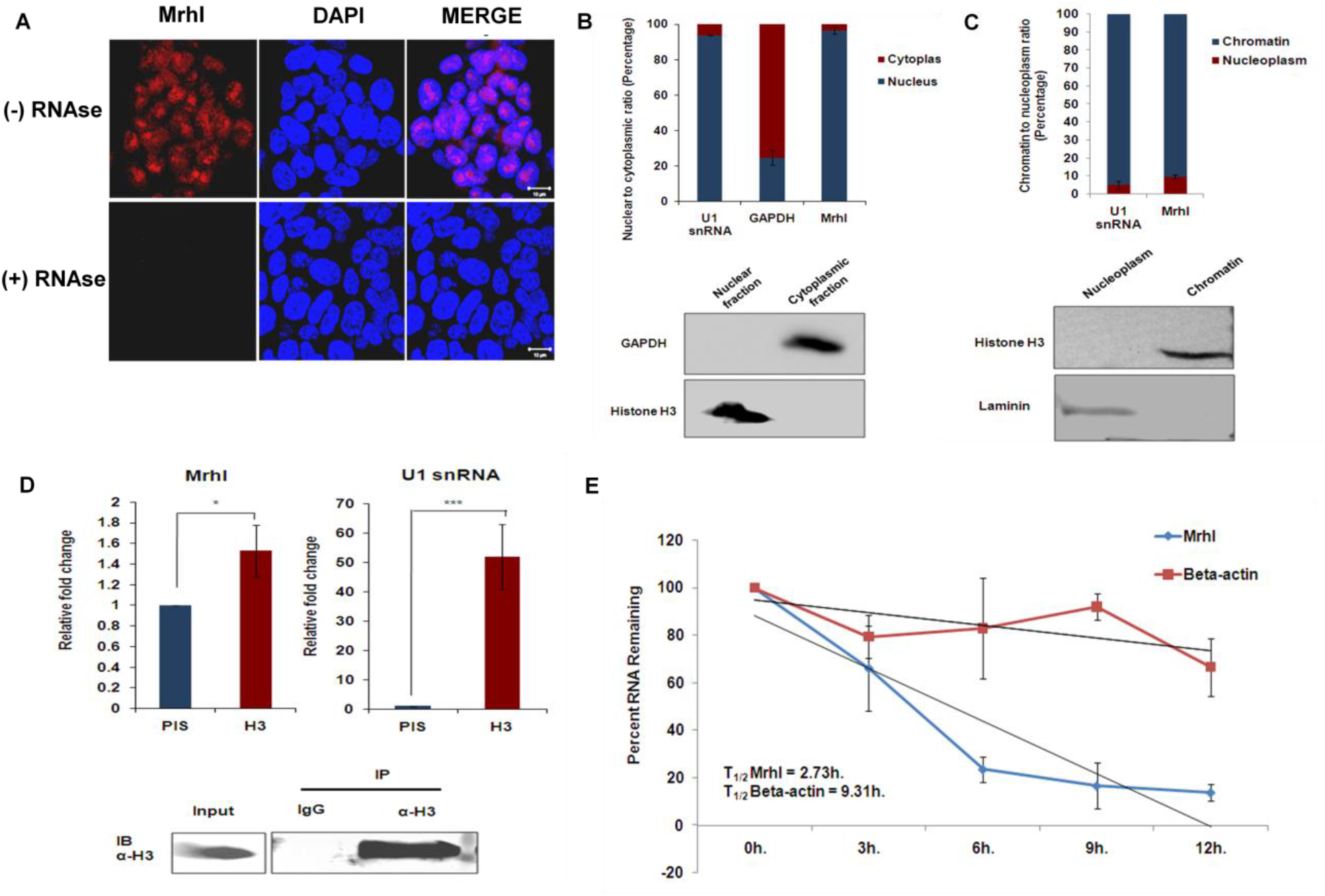
Mrhl is a nuclear-localized, chromatin bound moderately stable lncRNA in mESCs. (A) RNA FISH shows nuclear localization of Mrhl in mESCs. Scale bar = 10μm; (B) Fractionation validated observations in panel a. Western blot shows purity of fractions; (C) Chromatin and nucleoplasm fractions of nuclei show localization of Mrhl to chromatin. Western blot shows purity of fractions; (D) H3 ChIP and qPCR reveal Mrhl is bound with the chromatin in mESCs; (E) Actinomycin D half-life assay for Mrhl in mESCs. Error bars indicate standard deviation from three independent experiments. *p<0.05, **p<0.01, ***p<0.001, student’s t-test; Scale bar = 10 μm.

### Differentiation assay, knockdown and transcriptome analyses reveal Mrhl to regulate development and differentiation circuits in mESCs

We next differentiated the mESCs into embryoid bodies and very interestingly observed that Mrhl was preferentially up regulated at days 4 and 6 of embryoid body formation (Figure 2A). In perspective of the negative feedback regulation between Mrhl and WNT signaling in spermatogonial progenitors and of WNT signaling contributing to mESC physiology (Atlasi et al., 2013; Price et al., 2013; Sokol, 2011), we questioned whether Mrhl would function through similar mechanisms in this context as well. We performed shRNA mediated knockdown of Mrhl in mESCs and scored for its levels using RNA FISH followed by the status of β-CATENIN localization by IF. We observed that in cells where Mrhl was depleted with high efficiency, β-CATENIN was still localized at the membrane indicating non-activation of the WNT pathway (Figure 2B). Furthermore, p68 IP revealed that Mrhl does not interact with p68 in mESCs (Figure 2C). Keeping these observations in mind, we performed transient knockdown of Mrhl in mESCs using four independent constructs, two of which (referred to as sh. 1 and sh. 4 henceforth) showed us an average down regulation of 50% (Figure 2D) and subjected the scrambled (scr.) and sh.4 treated cells to analysis by RNA-Seq. A quick comparison of the FPKM values for Mrhl obtained in our analysis versus those reported in the ENCODE database as well as of scr. versus sh.4 displayed Mrhl to be a low abundant lncRNA and confirmed our knockdown efficiency respectively (**Figure 2-supplementary figure 2A**). Furthermore, we observed that the expression of pluripotency factors OCT4 (OCT), SOX2 and NANOG were not affected upon Mrhl knockdown in mESCs (**Figure 2-supplementary figure 2B**). We obtained a total of 1143 genes which were dysregulated in expression with 729 being down regulated and 414 being up regulated in expression (Figure 2E) and we refer to them as the differentially expressed genes (DEG, **Supplementary File 1**). The DEG majorly belonged to the class of protein-coding genes and transcription factors with an interesting proportion belonging to lincRNAs and antisense RNAs as well. Gene ontology (GO) analysis revealed diverse molecular functions such as binding (23.6%), catalytic activity (18.3%), receptor activity (8.8%) and signal transducer activity (7.4%) and biological processes such as cellular processes (40.9%), biological regulation (15.8%), metabolic process (25.7%), developmental process (11.7%) and multicellular organismal process (10.7%) to be affected (Figure 2F). We next performed a GO enrichment analysis with a p-value<0.05 to understand if one or more of the perturbed processes/pathways were statistically over represented over the others and we found positive regulation of developmental processes and positive regulation of multicellular organismal processes to be two such enriched perturbed processes (Figure 2G). Other processes such as protein metabolic processes, lipid metabolic processes, phosphate metabolic processes, protein modification processes and MAPK cascade were also amongst the GO enriched list of processes. We then performed Fisher’s exact test in the PANTHER interface with a p-value<0.001 and obtained several interesting GO categories to be further enriched and represented such as cell-cell signaling, ion transport, synaptic transmission, response to endogenous stimuli, ectoderm development, anion transport, cell-cell adhesion and neuromuscular synaptic transmission (**Supplementary Table 1**). The category of developmental processes (GO:0032502, **Supplementary File 2**) posed as the most interesting one since it has appeared in both the enrichment analyses and possessed the maximum number of perturbed genes i.e., 60 with a significant p-value of 6.44E-05. We examined the DEG belonging to this category and found that they belonged to two broad groups of lineage-specific TFs and cell adhesion/receptor activity. The former group comprised genes encoding factors involved in neuronal lineage, hematopoietic and vascular lineage, cardiac lineage, skeletal lineage, mesodermal lineage and pancreatic lineage (**Supplementary Table 2A**) whereas the latter group consisted of genes responsible for such functions as migration, axon guidance, signaling, growth and differentiation, structural roles and cellular proliferation amongst others (**Supplementary Table 2B**). From our assays and analyses herewith, we conclude that Mrhl majorly acts to regulate differentiation and development related genes and processes in mESCs. Also, we compared the DEG in Mrhl knockdown conditions in mESCs and GC1-Spg spermatogonial progenitors and made two observations: firstly, the perturbed transcriptome is vastly different, emphasizing the context-dependent role of Mrhl and secondly, about 25 genes were in common between the two datasets, suggesting some common target genes of regulation for Mrhl across developmental model systems (**Figure 2-supplementary figure 2C**).

**Figure 2:**
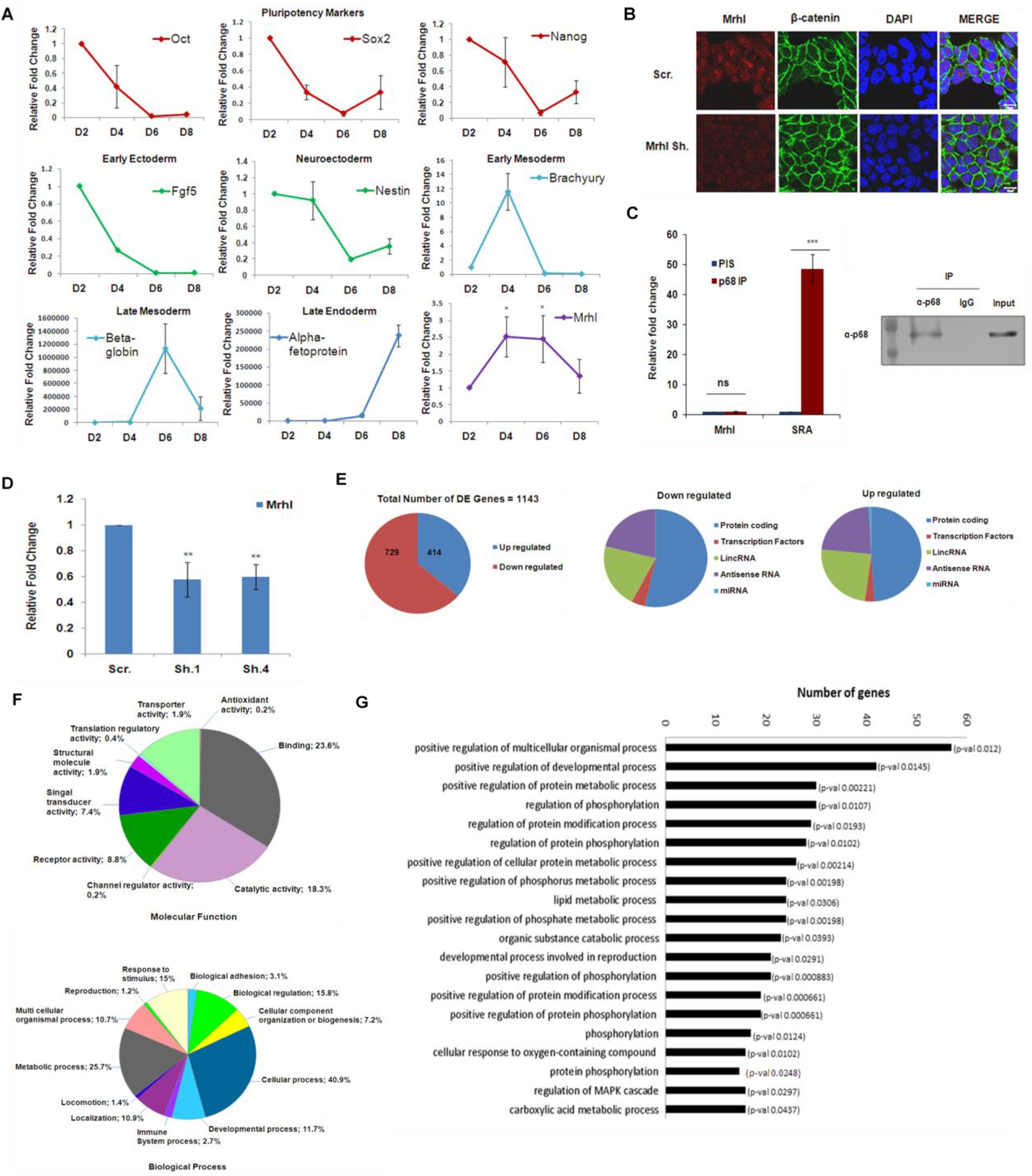
Analysis of role of Mrhl in mESCs through differentiation assay, knockdown and transcriptome analysis. (A) Mrhl shows differential expression during embryoid body differentiation of mESCs; (B,C) Mrhl does not function through the WNT/p68 cascade unlike in spermatogonial progenitors; (D) Knockdown efficiency of Mrhl in mESCs through two independent constructs i.e. sh.1 and sh.4 as compared to scrambled (scr.); (E) Representation of DEG classification; (F) Gene ontology analysis of DEG; (G) GO enrichment analysis of DEG. Error bars indicate standard deviation from three independent experiments. *p<0.05, **p<0.01, ***p<0.001, student’s t-test; Scale bar = 10 μm.

### Gene co-expression and TF interaction analyses show unique networks to be coordinated by Mrhl in mESCs

In order to organize the perturbed transcriptome under conditions of Mrhl knockdown in mESCs into functional and biologically relevant modules (W. Chen et al., 2018; X. Chen et al., 2016), we performed hierarchical clustering of the DEG and visualized the resultant clusters or modules with Cytoscape (Figure 3A). We obtained nine such co-expression modules. We then interrogated each of the modules for functional enrichment with GeneMania tool and observed diverse functions to be co-regulated by these groups of genes such as ion channel and transporter activities for clusters 4 and 5, nervous system functions for cluster 6, immune system processes for cluster 7 and responses to xenobiotic stimuli for cluster 8. Clusters 2 and 3 had varied functional enrichments whilst clusters 1 and 9 did not show any of such functional representations. These analyses further support our earlier observations all of which suggest that in mESCs, Mrhl acts to primarily modulate development and differentiation related processes.

**Figure 3:**
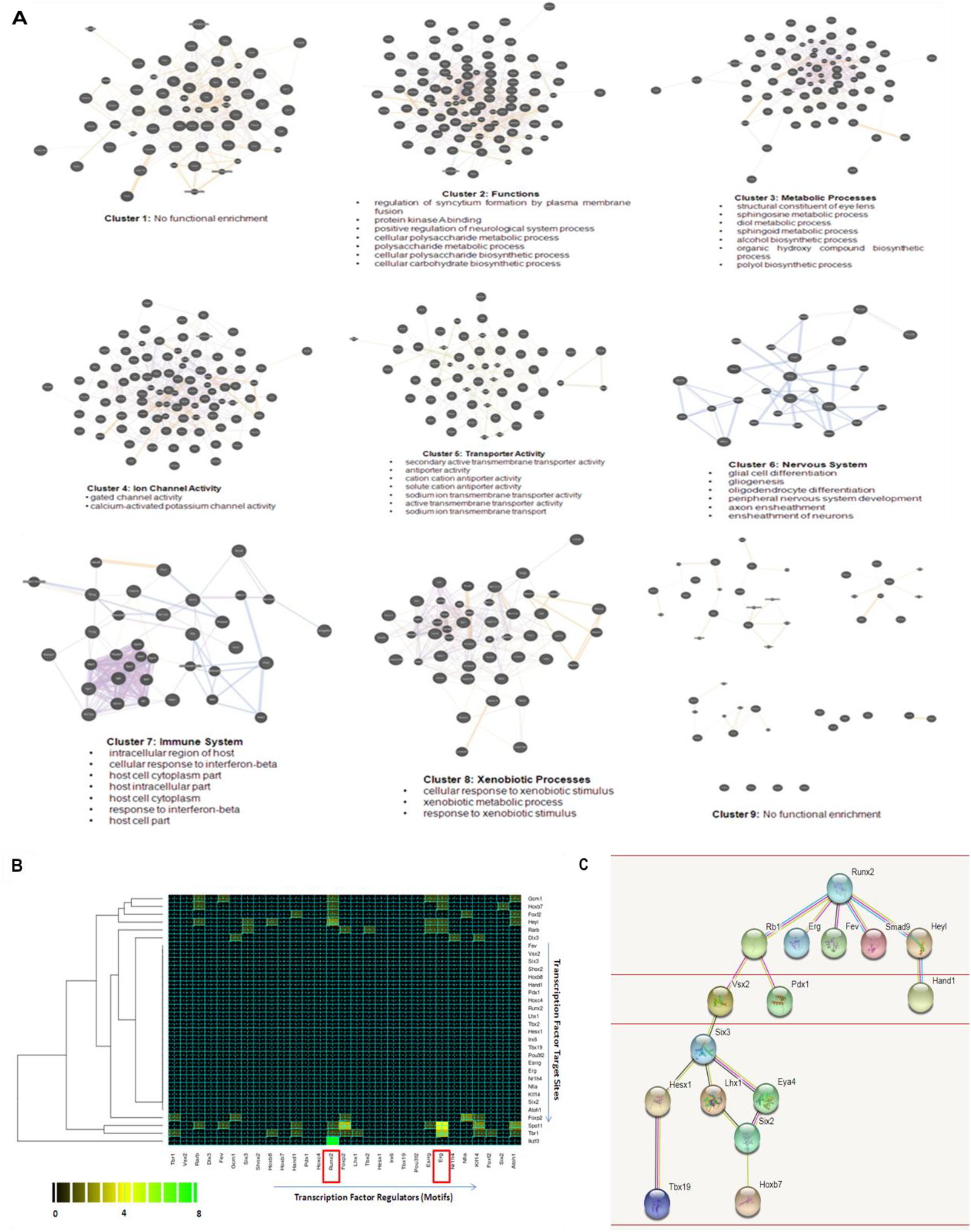
Gene co-expression and TF network analyses. (A) Gene co-expression modules and their corresponding functional enrichments; (B) Heat map visualization of TF matrix; (C) TF hierarchy as visualized in STRING.

We also performed a TF-TF interaction analysis to understand the potential cross-talk between the dysregulated set of TFs (**Figure 3-Supplementary File 3**) and to identify a master TF through which Mrhl might be acting to regulate the TF network. TFs have been implicated often in determining cellular states or fates (Dunn, Martello, Yordanov, Emmott, & Smith, 2014; Goode et al., 2016). A gene ontology analysis of the perturbed TFs revealed metabolic processes and developmental processes to be over represented in function (**Figure 3-supplementary figure 3A**). A heat map was generated for delineating the potential TF network by scanning the promoter sequence of each TF with the motif for each TF (Fig. 3B). RUNX2 was obtained as a potential master TF since it had the maximum number of motifs for binding across the promoters of all the other TFs followed by ERG (**Figure 3-supplementary figure 3B**). Based on this logic, a simplified TF hierarchy was established using STRING wherein we observed RUNX2 to be at the top of the hierarchy with RB1, ERG, FEV, SMAD9 and HEYL being in the first tier (Figure 3C) and undergoing regulation via RUNX2, although their direct regulation by Mrhl cannot be ruled out. Thus, we report a novel TF network or hierarchy operating in mESCs in the context of Mrhl. Furthermore, since all the TFs are involved in the specification of one of the three lineages, the observations herewith further emphasize on Mrhl acting to control cell fate specification and differentiation related circuits in mESCs.

### Stable knockdown of Mrhl in mESCs shows aberrance in lineage specification

Towards understanding the phenotypic implications of Mrhl depletion in mESCs and of our transcriptome analyses, we generated stable knockdown lines for Mrhl sh.4 and scr. control. Our initial characterization of the stable knockdown cells showed a knockdown efficiency of 50% (Figure 4A), no change in the pluripotency markers Oct 4, Sox2 and Nanog (Figure 4B,C and **Figure 4-supplementary figure 4A**) as well as no differences in the alkaline phosphatase expression levels (**Figure 4-supplementary figure 4B**) in the knockdown versus control lines. Since a couple of cell adhesion genes were present amongst the DEG, we performed IF for E-CADHERIN and F-ACTIN. A comparison of the staining intensities revealed no significant changes in cell adhesion properties between knockdown and control mESCs (Figure 4D and **Figure 4-supplementary figure 4C**). These observations corroborate with our transcriptome data. Next, keeping in mind our earlier conclusions and the observation that Mrhl is up regulated in expression during embryoid body differentiation, we subjected the knockdown and control cells to differentiation by the formation of embryoid bodies. Interestingly, we observed that over days 2 to 8 of differentiation, there was a marked aberrance in the specification of lineages, specifically the ectoderm and mesoderm lineages (Figure 4E, F). For the ectoderm lineage, there appears to be premature specification in the knockdown cells at day 2 with subsequent loss in maintenance of the lineage at days 6 and 8. The mesoderm lineage appears to be under-specified from the beginning of differentiation with some expression of the marker T at day 6 in the knockdown cells These observations imply that Mrhl is required in mESCs to undergo proper specification of early lineages, majorly the ectoderm and mesoderm lineages, although it might not have a specific role in mESCs per se in the absence of differentiation cues.

**Figure 4:**
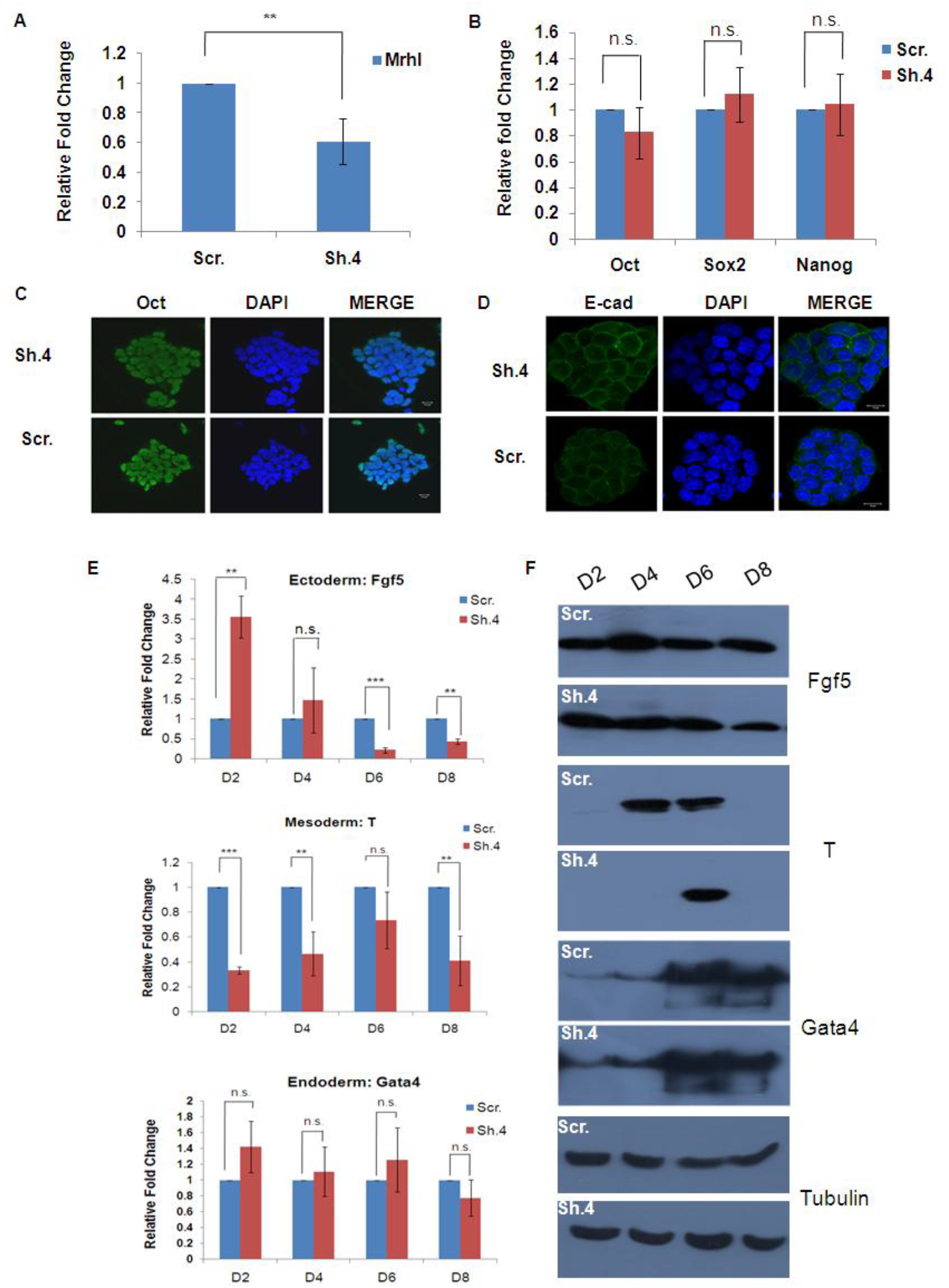
Stable knockdown of Mrhl in mESCs causes no change in pluripotency status but aberrance in lineage specification. (A) Knockdown efficiency in puromycin selected stable knockdown cells; (B) qRT-PCR for pluripotency markers; (C) Oct4 IF to confirm pluripotency status; (D) E-cadherin IF to check for cell adhesion status; (E) qRT-PCR for lineage-specific markers upon EB differentiation of stable knockdown cells as compared to scrambled control; (F) Western blots to confirm altered expression of lineage markers. Error bars indicate standard deviation from three independent experiments. *p<0.05, **p<0.01, ***p<0.001, student’s t-test; Panel E is representative data from one of three independent experiments, each carried out in biological triplicates; Scale bar

### Mrhl regulates target loci in mESCs through chromatin-mediated regulation

In order to further delineate the mechanism by which Mrhl regulates differentiation and development related genes and processes in mESCs, we performed genome-wide chromatin occupancy studies through ChIRP-Seq since we have demonstrated Mrhl to be a chromatin bound lncRNA in mESCs. We obtained a total of 21,997 raw peaks and after keeping a cutoff value of 5-fold enrichment, we obtained 21,282 peaks (Figure 5A, **Supplementary File 4**), emphasizing the significant and widespread chromatin occupancy of Mrhl in mESCs. The peak lengths and fold changes appear to be distributed equally across all the chromosomes (**Figure 5-supplementary figure 5A, B**). An annotation of the enriched peaks showed us that diverse regions including intronic, intergenic and promoter regions of genes as well as repeat elements undergo physical association with Mrhl (Figure 5B). Next, we overlapped the peaks from ChIRP-Seq and the DEG from RNA-Seq to understand what proportion of the dysregulated genes upon Mrhl knockdown is regulated by it at the chromatin level. In this regard, we have used −10 kb upstream of transcription start site (TSS) and +5 kb downstream of TSS of genes as our domain of target gene regulation by Mrhl which narrowed down the number of peaks to 3412. The overlap analysis revealed 71 genes which are physically occupied and are also regulated by Mrhl at the chromatin level (Figure 5C, **Supplementary File 5**). We further examined the 71 genes in detail and found a subset of six genes i.e., RUNX2, SIX2, DLX3, HOXB7, POU3F2 and FOXP2 to be of noticeable important in terms of being firstly, lineage determining or development associated transcription factors and secondly, being physically occupied by Mrhl at the promoter (**Supplementary Table 3A**).

**Figure 5:**
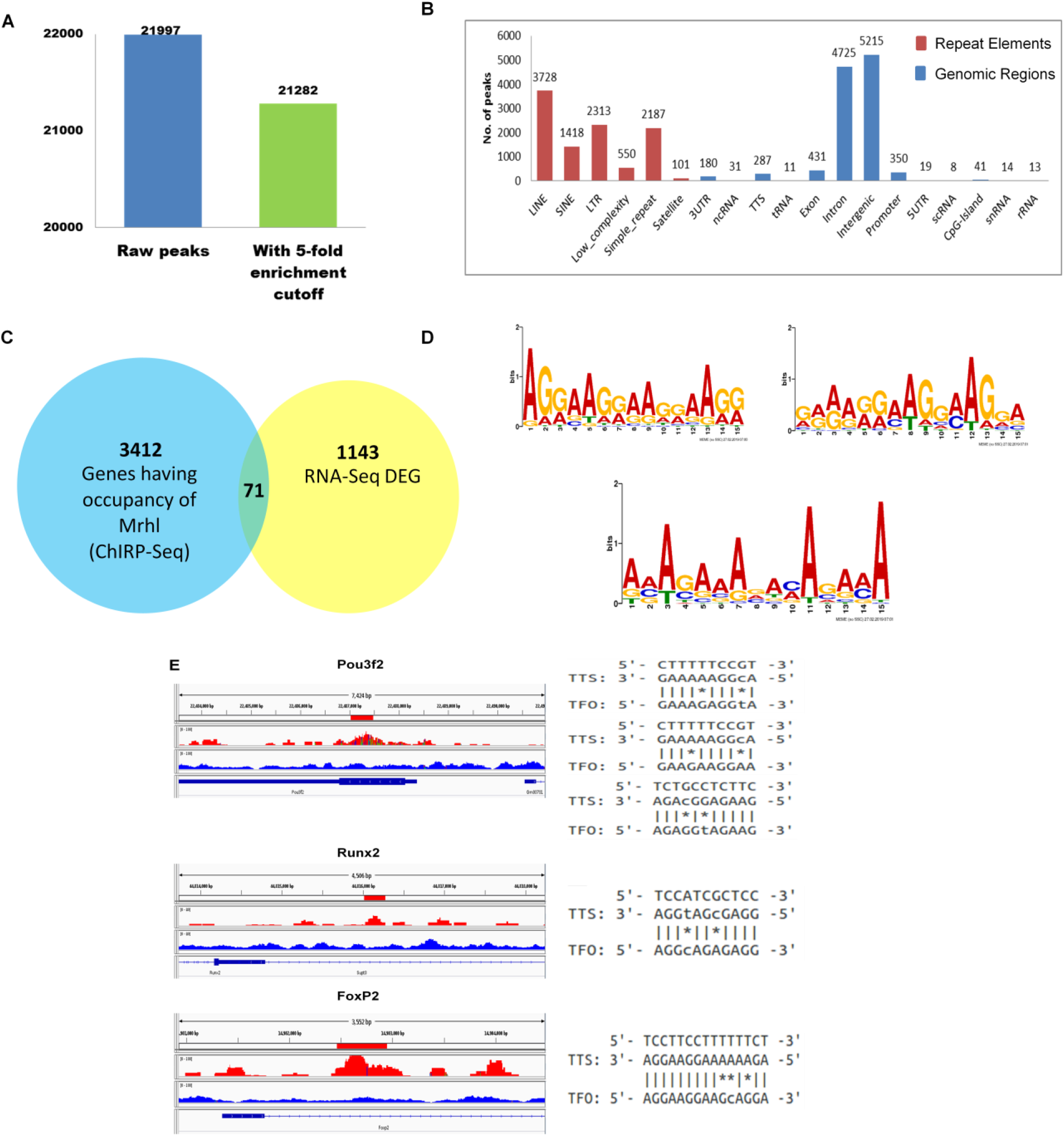
ChiRP-Seq analysis for Mrhl in mESCs. (A) Number of peaks obtained before and after cutoff; (B) Annotation of peaks; (C) Overlap of ChIRP-Seq and RNA-Seq datasets; (D) Motif analysis for genome occupancy for Mrhl on target genes; (E) Triplex formation prediction at select loci involved in development and differentiation functions.

A recently established mechanism of chromatin-mediated target gene regulation by lncRNAs is via the formation of RNA-DNA-DNA triple helical structures (Mondal et al., 2015; Postepska-Igielska et al., 2015; S. Wang et al., 2018). To delve deeper into the exact mechanism of chromatin-mediated gene regulation by Mrhl at the promoter of the aforesaid developmental loci, we hypothesized triplex formation by Mrhl at these loci. For this, we selected a subset of the six genes i.e., RUNX2, HOXB7, FOXP2 and POU3F2. Their roles in governing the development of specific lineages such as the osteoblast lineage [RUNX2 (Komori, 2002)], neuronal lineage and brain development [FOXP2 (Chiu et al., 2014; Tsui, Vessey, Tomita, Kaplan, & Miller, 2013), POU3F2 (Y. J. Lin et al., 2018; Urban et al., 2015)) or having multiple functions during development [HOXB7 (Candini et al., 2015; Klein, Benchellal, Kleff, Jakob, & Ergun, 2013)] have been widely established. We first performed a search for motifs for binding to target chromatin loci by Mrhl which led to the identification of three distinct motifs with motif 1 being present in 21.46%, motif 2 being present in 28.16% and motif 3 being present in 43.08% of the total number of peaks. (Figure 5D and **supplementary table 3B**). Next, we analyzed the promoters of the aforementioned genes corresponding to the region where ChIRP-Seq peaks were obtained for the presence of one or more of the three motifs by FIMO in MEME suite followed by potential triplex formation at those sites using the Triplexator program. We observed the presence of only one motif (motif 3) at the Mrhl occupied region in POU3F2, motifs 1, 2 and 3 in FOXP2 and again motif 3 in RUNX2 and HOXB7. Interestingly, triplex prediction revealed triplex forming potential at two different sites in POU3F2, one lying within the motif sequence whereas in FOXP2 and RUNX2, potential triplex forming sites were found immediately adjacent to the motif sequences. HOXB7, however, did not show propensity for triplex formation within the Mrhl occupied region (Figure 5E, **Supplementary table 3C, Supplementary File 6**). In lieu of these observations, we conclude that Mrhl regulates key lineage-specific TFs at the chromatin possibly through triple helix formation to regulate differentiation of mESCs.

## Discussion

Regulation of pluripotency and/or differentiation phenomena in embryonic stem cells by lncRNAs in combination with their extensive context-dependent roles poses them as novel therapeutic targets in the context of regenerative medicine. Linc-RoR mediates the formation of human induced pluripotent stem cells (Loewer et al., 2010) and contributes to human embryonic stem cell self-renewal (Y. Wang et al., 2013) whereas lncRNA Cyrano is involved in maintenance of pluripotency of mESCs (Smith, Starmer, Miller, Sethupathy, & Magnuson, 2017). In parallel, linc-RoR has been implicated in osteogenic differentiation of mesenchymal stem cells (Feng et al., 2018) whilst Cyrano has been shown to function in conjunction with other non-coding RNAs to regulate neuronal activity in the mammalian brain (Kleaveland, Shi, Stefano, & Bartel, 2018) as well as neurodevelopment in zebrafish (Sarangdhar, Chaubey, Srikakulam, & Pillai, 2018). In our current study, we have characterized and addressed the functional implications of lncRNA Mrhl, previously shown to regulate male germ cell meiotic commitment, in mESC circuitry to understand its role in development and differentiation. The role of nuclear lncRNAs in coordinating and controlling gene expression at a genome-wide level through a myriad of mechanisms is well known, amongst which regulation of chromatin architecture/chromatin state at target loci is predominant (Ballarino et al., 2018; Cajigas et al., 2015; Sun, Hao, & Prasanth, 2018).

LncRNA Mrhl depletion majorly dysregulates pathways and processes in mESCs which are related to lineage-specific development and differentiation and which are largely distinct from the perturbed gene set in spermatogonial progenitors. This emphasizes the context-dependant role of Mrhl as a molecular player of the cellular system. This is further supported by lack of association of Mrhl with p68 or absence of regulation of WNT signaling in mESCs as we have shown in our studies. In perspective, it would be interesting to address the protein interaction partners of Mrhl in mESCs, especially to understand in greater depth how Mrhl mediates regulation at target genes.

An intriguing point to note is absolutely no perturbation of pluripotency status of mESCs upon Mrhl knockdown rather disruption of lineage specification in embryoid bodies. These observations suggest Mrhl to be involved in specifying a primed state of the mESCs wherein they can undergo balanced specification of lineages upon obtaining differentiation cues. Other disrupted pathways such as ion transport which have been recurrent in all systems analyses would be an interesting aspect to address in the future. The aforesaid conclusions are further supported by the fact that Mrhl is up regulated in expression at days 4 and 6 during embryoid body differentiation, suggesting it as a regulator of differentiation programs in mESCs and explaining the aberrance in lineage specification upon differentiation of Mrhl knockdown stable mESCs. The lack of any effect on the endoderm lineage during in vitro differentiation of the stable knockdown cells is explainable from an analysis of GO:0032502 wherein in vast majority only ectoderm and mesoderm related genes were observed to be dysregulated. Furthermore, in our ENCODE data analysis of organs, Mrhl was observed to be expressed predominantly in embryonic stages of organs of various lineages such as the brain (ectoderm), kidney, testes (mesoderm) and lung, liver (endoderm). An interrogation of Mrhl expression in the recently released Mouse Organogenesis Cell Atlas (Cao et al., 2019) showed Mrhl to be expressed in progenitor cell types of various tissues. This can only give us an insight about the involvement of Mrhl not only in the early stages of germ layer specification but in the later stages of organogenesis as well, although the exact functions in the latter need to be still addressed.

The regulation of a novel TF network in mESCs by Mrhl comprising mostly lineage-determining TFs is an exciting observation. TFs act to govern gene expression programs defining particular cellular states, more so in association with other TFs in a network (Dore & Crispino, 2011; Niwa, 2018). LncRNAs often integrate into such networks by regulating TFs individually or via a master TF(s) resulting in downstream regulation of gene expression (Herriges et al., 2014; Yo & Runger, 2018). The TF network operating in mESCs under the regulation of Mrhl has RUNX2 at the top of the hierarchy posing it as a master TF in the hierarchy and being potentially regulated by Mrhl directly through triplex formation at the chromatin level. Amongst the 71 genes overlapping between the DEG and the ChIRP-Seq reads, RUNX2, SIX2 and HOXB7 belong to a common group of the aforesaid set and the TF network. The absence of triplex formation sites in HOXB7 in spite of the presence of Mrhl binding motifs in its promoter further strengthens the hypothesis that Mrhl might be regulating the entire network through RUNX2. Additionally, FOXP2 and POU3F2 although not a part of the TF network, display triplex forming potential within the Mrhl occupied regions. These observations suggest an interesting mechanism wherein Mrhl regulates differentiation programs in mESCs through direct regulation of relevant TFs at their promoters.

A further analysis of our transcriptome studies herewith shows a significant over representation of dysregulated genes and processes belonging to neuronal lineage. Ectoderm development was one of the enriched processes in the Fisher’s exact test and ~20% of the dysregulated genes in GO: 0032502 belonged to neuronal lineage development related functions. In the gene co-expression analysis, nervous system emerged as one of the perturbed clusters. Furthermore, Mrhl is predicted to regulate an important neuronal TF such as POU3F2 directly at the chromatin level. Whilst our phenotype analysis of the stable knockdown cells showed perturbations in early specification of the ectoderm and mesoderm lineages, further investigations of the role of Mrhl in specifying more specialized lineages such as the neuronal lineage would be an interesting aspect of study.

LncRNAs have been implicated widely for their contributions to embryonic stem cell differentiation, cell fate specification, organogenesis and development through a diverse array of mechanisms (Grote & Herrmann, 2015; Perry & Ulitsky, 2016; Sarropoulos, Marin, Cardoso-Moreira, & Kaessmann, 2019). In our studies we have characterized lncRNA Mrhl and its functional significance in mESC towards decoding its role in developments. We show Mrhl to regulate downstream genes and processes involved in differentiation and lineage specification that was reflected in our phenotype studies. A major finding of this study was its potential direct chromatin mediated regulation of key TFs that mediate differentiation of stem cells into a specific lineage. Overall, we establish lncRNA Mrhl to be a mediator of differentiation and cell fate specification events in mESCs.

## Materials and Methods

### Cell lines, antibodies, plasmids and chemicals

E14tg2a feeder independent mESC line was a kind gift from Prof. Tapas K. Kundu’s lab (JNCASR, India). mESCs were maintained on 0.2% gelatin coated dishes in mESC medium containing DMEM, high glucose, 15% FBS, 1X non-essential amino acids, 0.1 mM β-mercaptoethanol and 1X penicillin-streptomycin. HEK293T cells (ATCC, U.S.A) were maintained in DMEM, 10% FBS and 1X penicillin-streptomycin.

The antibodies used in this study are as follows: Anti-GAPDH (Abeomics, ABM22C5), anti-H3 (Abcam, ab46765), anti β-CATENIN (Abcam), anti p68 (Novus Biologicals), anti-OCT4 (Abcam, ab27985).

Scrambled and Mrhl shRNA plasmids 1, 2, 3 and 4 were custom made from Sigma in the pLKO.1-Puro-CMV-tGFP vector backbone. The sequences of the shRNAs are as follows: Mrhl shRNA#1: 5’GCACATACATACATACACATATATT 3’, Mrhl shRNA#4: 5’GGAGAAACCCTCAAAAGTATT 3’.

All fine chemicals were obtained from Sigma (unless otherwise mentioned), gelatin was obtained from Himedia and FBS was obtained from Gibco (Performance Plus, US Origin).

### Protocols for mESC differentiation, transfection and cell assays

For embryoid body (EB) differentiation, E14tg2a cells were trypsinized using 0.05% trypsin and 2.5×10^5^ cells were plated onto 35 mm bacteriological grade dishes (Tarsons) in EB differentiation medium containing DMEM, 10% FBS, 0.1 mM β-mercaptoethanol and 1X penicillin-streptomycin. EBs were harvested at different time points and processed accordingly for further analysis.

All transfections for E14tg2a cells were performed using Trans IT-X2 reagent (Mirus, MIR 6003) as per the manufacturer’s protocol. For transient knockdown analysis, cells were harvested 48 hours post-transfection. Transfections for HEK293T cells were performed using Lipofectamine 2000 (Thermo Fisher Scientific) as per the manufacturer’s protocol.

For measurement of half-life of Mrhl, E14tg2a cells, at a confluency of ~80-90%, were treated with 10 μM actinomycin D (Sigma, A9415). Cells were harvested at different time points for further analysis.

Alkaline phosphatase assay was performed as per the manufacturer’s protocol (Sigma, 86R).

### Generation of stable knockdown lines

Stable mESC knockdown lines for Mrhl were generated in E14tg2a cells as per the protocol of Pijnappel *et al* (Pijnappel, P.A. Baltissen, & Timmers, 2013) with some modifications Briefly, viral particles were generated in HEK293T cells by transfection of 5 μg scrambled or Mrhl shRNA plasmids, 2.5 μg pSPAX2, 1.75 μg pVSVG and 0.75μg pRev. The media containing viral particles was harvested 48 hours after transfection and centrifuged at 4,000 rpm for 5 minutes to remove cellular debris. Fresh mESC medium was added and harvested after an additional 24 hours to collect second round of viral particles. The viral supernatants were stored in −80°C if necessary. E14tg2a cells were plated such that they reached a confluency of ~60-70% on the day of transduction. The viral supernatant was mixed with 8 μg/ml DEAE-dextran and 1000 units/ml ESGRO (Merck Millipore) and added directly to the E14tg2a cells. Transduction was performed for 24 hours with the first round of viral particles and an additional 24 hours with the second round of viral particles. The transduced cells were then subjected to puromycin selection (1.5 μg/ml puromycin) for a week.

### RNA fluorescent in situ hybridization (FISH) and immunofluorescence (IF)

RNA FISH followed by IF was performed as per the protocol of de Planell-Saguer *et al* (de Planell-Saguer, Rodicio, & Mourelatos, 2010) with modifications. The probes used for RNA FISH studies were Cy5 labelled locked nucleic acid probes procured from Exiqon (reported in (Arun et al., 2012)). Briefly, cells were grown on gelatin coated coverslips. EBs were fixed in 4% paraformaldehyde for 7-8 hours at 4°C followed by equilibration in 20% sucrose solution overnight and embedded in tissue freezing medium (Leica). The embedded tissues were then cryo-sectioned and collected on Superfrost slides (Fisher Scientific). For cells, a brief wash was given with 1X PBS (phosphate buffered saline, pH 7.4) followed by fixation with 2% formaldehyde for 10 minutes at room temperature. The cells were then washed with 1X PBS three times for 1 minute each and permeabilization buffer (1X PBS, 0.5% Triton X-100) was added for 5 minutes and incubated at 4°C. The permeabilization buffer was removed and cells were washed briefly with 1X PBS for three times at room temperature. For tissue sections, antigen retrieval was performed by boiling the sections in 0.01M citrate buffer (ph 6) for 10 minutes. The sections were allowed to cool, washed in distilled water three times for 5 minutes each and then in 1X PBS for 5 minutes, each time with gentle shaking.

For FISH, samples were blocked in prehybridization buffer [3% BSA, 4X SSC (saline sodium citrate, pH 7)] for 40 minutes at 50°C. Hybridization (Mrhl probes tagged with Cy5, final concentration 95nM) was performed with prewarmed hybridization buffer (10% dextran sulphate in 4X SSC) for 1 hour at 50°C. After hybridization, slides were washed four times for 6 minutes each with wash buffer I (4X SSC, 0.1% Tween-20) at 50°C followed by two washes with wash buffer II (2X SSC) for 6 minutes each at 50°C. The samples were then washed with wash buffer III (1X SSC) once for 5 minutes at 50°C followed by one wash with 1X PBS at room temperature. For tissue sections, all washes were performed as mentioned above with a time of 4 minutes for buffers I-III.

For IF, samples were blocked with IF blocking buffer (4% BSA, 1X PBS) for 1 hour at 4°C. Primary antibody solution (2% BSA, 1X PBS) was prepared containing the appropriate dilution of desired primary antibody and the samples were incubated in it for 12 hours at 4°C. Next day, the samples were washed with IF wash buffer (0.2% BSA, 1X PBS) three times for 5 minutes each with gentle shaking. The samples were incubated in secondary antibody for 45 minutes at room temperature and washed with 1X PBS three times for 10 minutes each with gentle shaking. The samples were finally mounted in mounting medium containing glycerol and DAPI.

### Biochemical fractionation

#### Cell fractionation

Approximately 5-10 million cells were lysed using lysis buffer (0.8 M sucrose, 150 mM KCl, 5 mM MgCl_2_, 6 mM β-mercaptoethanol and 0.5% NP-40) supplemented with 75 units/ml RNAse inhibitor (Thermo Fisher Scientific) and 1X mammalian protease inhibitor cocktail (Roche) and centrifuged at 10,000 g for 5 minutes at 4 °C. The supernatant containing cytoplasmic fraction was mixed with 3 volumes of TRIzol for RNA extraction or with Laemmli buffer for western blotting as described later. The resultant pellet was washed twice with lysis buffer and was subjected to RNA or protein extraction.

#### Sub-nuclear fractionation

Approximately 10 million cells were lysed with hypotonic lysis buffer (10 mM Tris-HCl ph 7.5, 10 mM NaCl, 3 mM MgCl_2_, 0.3% v/v NP-40 and 10% v/v/ glycerol) supplemented with RNAse inhibitor and mammalian protease inhibitor cocktail and centrifuged at 1,000 g for 5 minutes at 4 °C. The supernatant comprising the cytoplasmic fraction was kept aside and the nuclear pellet was washed twice with hypotonic lysis buffer. The nuclear pellet was then resuspended in modified Wuarin-Schibler buffer (10 mM Tris-HCl pH 7.0, 4 mM EDTA, 300 mM NaCl, 1 M urea and 1% NP-40) supplemented with RNAse inhibitor and mammalian protease inhibitor cocktail and vortexed for 10 minutes. Nucleoplasmic and chromatin fractions were separated by centrifugation at 1,000 g for 5 minutes at 4 °C. The chromatin pellet was resuspended in sonication buffer (20 mM Tris-HCl pH 7.5, 150 mM NaCl, 3 mM MgCl_2_, 0.5 mM PMSF, 75 units/ml RNAse inhibitor) and sonicated for 10 minutes. The chromatin was then obtained as the supernatant following centrifugation at 18,000 g for 10 minutes at 4 °C to remove all debris. The resultant nucleoplasmic and chromatin fractions were then subjected to RNA or protein extraction as described later.

### Immunoprecipitation (IP)

#### p68 IP

Cells were lysed in hypotonic lysis buffer (10 mM Tris-HCl ph 7.5, 10 mM NaCl, 3 mM MgCl_2_, 0.3% NP-40, 10% glycerol) supplemented with RNase inhibitor, mammalian protease inhibitor cocktail and 1mM PMSF. Nuclei were pelleted down at 1200 g for 10 minutes at 4°C and subsequently lysed in nuclear lysis buffer (150 mM KCl, 25 mM Tris pH 7.4, 5 mM EDTA, 0.5% NP-40) supplemented with RNase inhibitor, mammalian protease inhibitor cocktail and PMSF. The debris was removed by centrifugation at 15,000 g for 10 minutes at 4°C and the supernatant nuclear fraction was collected. To 1 mg of the nuclear fraction containing proteins, 7 μg of either pre-immune serum or p68 antibody was added and incubated overnight at 4°C. Next day, the fraction was incubated with protein A dynabeads for 3 hours at 4°C. The beads were washed with wash buffer (20 mM Tris-HCl pH 7.4, 2 mM MgCl_2_, 10 mM KCL, 150 mM NaCl, 10% glycerol, 0.2% NP-40) supplemented with RNase inhibitor, mammalian protease inhibitor cocktail and PMSF. Subsequently, the beads were washed twice with wash buffer (as above with 0.5% NP-40) and collected. The beads were then resuspended either in RNAiso Plus and subjected to RNA isolation for qRT-PCR analysis or in Laemmli buffer and resolved on a 10% SDS-PAGE gel for western blotting analysis as described later.

#### Chromatin IP

Chromatin IP (ChIP) was performed as per Cotney and Noonan’s protocol (Cotney & Noonan, 2015). Approximately 6-7 million cells were cross-linked with 1% formaldehyde for 10 minutes. The reaction was quenched with 125mM glycine for 5 minutes and washed twice with ice-cold 1X PBS. Cells were lysed using hypotonic lysis buffer (10mM Tris ph 7.5, 10mM NaCl, 3mM MgCl_2_, 0.3% NP-40 and 10% glycerol) supplemented with RNAse inhibitor and mammalian protease inhibitor cocktail. Nuclei were pelleted at 1200g for 10 minutes at 4°C and resuspended in nuclear lysis buffer (0.1% SDS, 0.5% Triton X-100, 20mM Tris pH 7.5 and 150mM NaCl) RNAse inhibitor and mammalian protease inhibitor cocktail. Resuspended nuclei were then sonicated (Biorupter, 25 cycles) and ~15 ug chromatin was incubated with either 4μg H3 antibody or pre-immune serum overnight at 4°C. Antibody bound chromatin was then pulled down using protein A dynabeads (Thermo Fischer Scientific) for 3 hours at 4°C. The beads were then washed sequentially with wash buffer I (nuclear lysis buffer), wash buffer II (nuclear lysis buffer with 500mM NaCl) and wash buffer III (10mM Tris pH 8.0, 0.5% NP-40, 0.5% sodium deoxycholate, 1mM EDTA), each supplemented with RNAse inhibitor and mammalian protease inhibitor cocktail for 5 minutes each. The beads were then either processed directly for western blotting or subjected to elution for 1 hour at 55°C in elution buffer (100mM NaCl, 10mM Tris pH 7.5, 1mM EDTA, 0.5% SDS) supplemented with 100ug/ml proteinase K. The supernatant was then subjected to reverse cross-linking by heating for 10 minutes at 95°C and then taken forward for RNA isolation using TriZol (Ambion).

#### Chromatin Isolation by RNA Purification (ChIRP)

ChIRP was carried out according to the protocol of Chu *et al* (Chu, Quinn, & Chang, 2012). Anti-sense (As) DNA probes with BiotinTEG at 3’ end was designed using the online probe designer at single-molecule FISH online designer. As-DNA probes were designed for both LncRNA Mrhl and LacZ for selective retrieval of RNA target by ChIRP. Cells were grown to confluency and ~80 million cells were harvested to perform ChIRP. Cells were cross linked with 1% glutaraldehyde to preserve RNA-Chromatin interactions and cell pellet was prepared. Crosslinked cells were lysed to prepare cell lysate using freshly prepared lysis buffer (50 mM Tris–Cl pH 7.0, 10 mM EDTA, 1 % SDS) supplemented with 1mM PMSF, 1X mammalian protease inhibitor cocktail (PI) and superase-in. The suspension was divided into 400 μl aliquots and subjected immediately to sonication. Sonication was continued until the cell lysate was no longer turbid. When lysate turned clear, 5 μl lysate was transferred to a fresh eppendorf tube. 90 μl DNA Proteinase K (PK) buffer (100 mM NaCl, 10 mM Tris–Cl pH 7.0, 1 mM EDTA, 0.5 % SDS) supplemented with 5 μl PK was added and incubated for 45 min at 55°C. DNA was extracted with Diagenode microChIP DiaPure purification kit to check DNA size on 1% agarose gel. If bulk of the DNA smear was 100-500 bp, sonication was assumed completed. Subsequent morning biotinylated DNA probes were hybridized to RNA and bound chromatin was isolated. For a typical ChIRP sample using 1 ml of lysate, 5% was kept aside for RNA input and DNA input. Hybridization buffer (750 mM NaCl, 1 % SDS, 50 mM Tris–Cl pH 7.0, 1 mM EDTA, 15 % formamide) supplemented with 1mM PMSF, 1X PI and superase-in was added to each lysate tube. 100 pmol of probe mix was added per 1 ml chromatin (1 μl of 100 pmol/μl probe per 1 ml chromatin). This was incubated for 4 hrs at 37 °C with shaking. 100 μl C-1 magnetic beads were used per 100 pmol of probes. Beads were resuspended in original volume of lysis buffer supplemented with PMSF, PI and superase-in. After 4 hr hybridization reaction was completed, 60 μl beads were added to each tube and incubated at 37 °C for 30 min with shaking. The beads were then washed for five times with wash buffer (2× SSC, 0.5 % SDS, 1 mM PMSF). At last wash, the beads were resuspended well. 100 μl was set aside for RNA isolation and 900 μl for DNA isolation. All tubes were placed on DynaMag-2 magnetic strip and last traces of wash buffer removed. RNA was isolated for further qRT-PCR analysis. LacZ was used as a negative control. DNA was isolated and sent for high throughput sequencing in duplicates.

#### RNA isolation and PCR

Total RNA was isolated from cells or tissues using TRIzol (Thermo Fisher Scientific) for RNA-sequencing and IP or using RNAiso Plus (Takara Bio) for analysis by qRT-PCR as per the manufacturer’s protocol. All RNA samples were subjected to DNase treatment (NEB) and about 1-3.5 μg of the RNA was taken for cDNA synthesis using oligodT primers (Thermo Fisher Scientific), RevertAid reverse transcriptase (Thermo Fisher Scientific) and RNase inhibitor (Takara Bio). The cDNA was diluted 1:1 with nuclease free water and subjected to qRT-PCR using SyBr green mix (Takara) in real-time PCR machine (BioRad CFX96). All semi-quantitative PCR was performed in thermal cycler machine (BioRad, Tetrad2) using Taq polymerase (Takara).

### Systems analysis

#### RNA-Seq analysis

E14tg2a cells treated with scrambled or Mrhl shRNA (shRNA 4) were subjected to RNA isolation and quality check. RNA samples were then subjected to library preparation in duplicates and sequenced on Illumina Hi-Seq 2500 platform. mm10 Genome was downloaded from GENCODE and indexed using Bowtie2-build with default parameters. Adapter ligation was done using Trim Galore (v 0.4.4) and each of the raw Fastq files were passed through a quality check using the FastQC. PCR duplicates were removed using the Samtools 1.3.1 with the help of ‘rmdup’ option. Each of the raw files were then aligned to mm10 genome assembly using TopHat with default parameters for paired-end sequencing as described in (Trapnell et al., 2012). After aligning, quantification of transcripts was performed using Cufflinks and then Cuffmerge was used to create merged transcriptome annotation. Finally differentially expressed genes were identified using Cuffdiff. The threshold for DE genes was log2 (fold change) >1.5 for up regulated genes and log2 (fold change) <1.5 for down regulated genes. The DE genes were analyzed further using R CummeRbund package.

#### GO enrichment analysis

Gene Ontology (GO) analysis was performed in PANTHER (Thomas et al., 2003). Significant enrichment test was performed with the set of differentially expressed genes in PANTHER and Bonferroni correction method was applied to get the best result of significantly enriched biological processes.

#### Fisher’s exact test

Fisher’s exact test was performed in PANTHER Gene Ontology (GO) where p-value significance was calculated based on the ratio of obtained number of genes to the expected number of genes (O/E) considering the total number of genes for the respective pathway in *Mus musculus*.

#### Cluster analysis

Hierarchical clustering method was performed using Cluster 3.0 (de Hoon, Imoto, Nolan, & Miyano, 2004). Gene expression data (FPKM of all samples i.e, scrambled and shRNA treated) was taken and log2 transformed. Low expressed (FPKM<0.05) and invariant genes were removed. Then genes were centered and clustering was performed based on differential expression pattern of genes and fold change. Genes were grouped in 9 clusters and visualized as a network in Cytoscape (Shannon et al., 2003). Functional enrichment of each cluster was performed using the Gene Mania Tool (Warde-Farley et al., 2010).

#### TF network analysis

Motifs were downloaded for all transcription factors from JASPAR (Mathelier et al., 2014) and sequence of interest for each TF (1.5 kb upstream & 500bp downstream of TSS) was extracted using BedtoFasta of the Bedtools suite (Quinlan & Hall, 2010). Then each motif was scanned across the sequence of all TFs to create the table matrix that reflects the number of binding sites for each TF across the other TFs using MEME suite (Bailey et al., 2009) with a e-value of 1E-04. Finally the heatmap was generated from the table matrix using R 3.3.2. TFs were fed into STRING (Jensen et al., 2009) to visualize the known interactions from the experimental data and hierarchy was setup manually as per the interaction among given TFs (proteins).

#### ChIRP-Seq analysis

mm10 Genome was downloaded from GENCODE and indexed using Bowtie2-build with default parameters. Adapter ligation was done using Trim Galore (v 0.4.4) and each of the raw Fastq files were passed through a quality check using the FastQC. PCR duplicates were removed using the Samtools 1.3.1 with the help of ‘rmdup’ option. Each of the raw files was then aligned to mm10 genome assembly using Bowtie 2 with default parameters for paired-end sequencing as described in. Replicates of both control and treated were merged respectively. Peaks were called using MACS2. Final peaks were selected giving the criteria of above 5-fold change and p value < 0.05.

#### Mrhl Motif Prediction

Motifs were identified using MEME, using the criteria of One Occurrence Per Sequence (OOPS) and significance of 1E-04 for 21282 genomic loci. Sequence for each locus was extracted from mm10 genome using bedtofasta of bedtools suite. After feeding sequences from 21282 genomic loci obtained from MACS2, 3 significant motifs were identified.

#### Triplex Prediction

Sequence from the Mrhl occupied region (in addition extended upto +/-25 bp) of selected genes was used for Triplex prediction using the software Triplexator (Buske, Bauer, Mattick, & Bailey, 2012) with default parameters.

#### qRT-PCR primer sequences

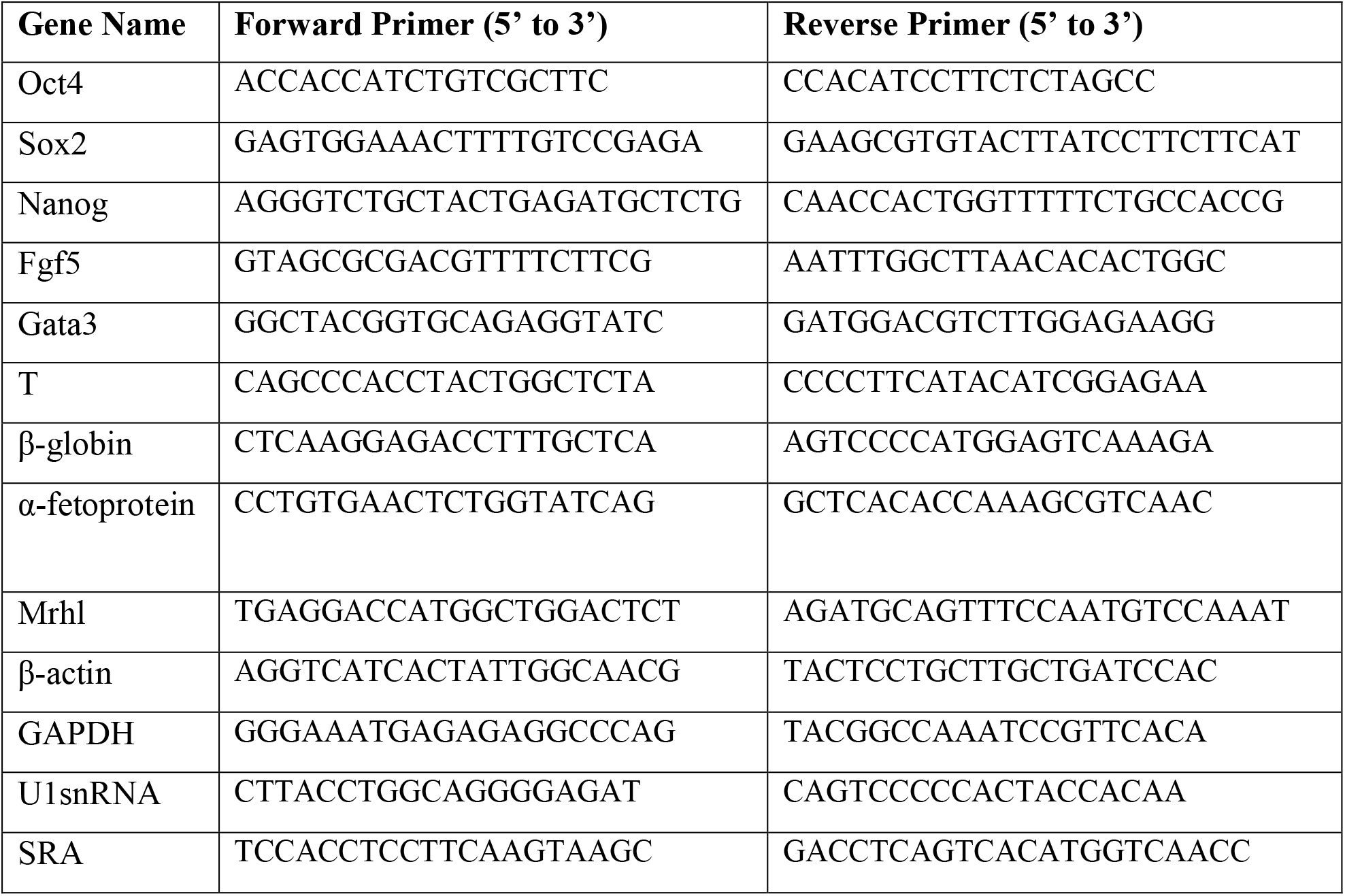

#### Oligo Sequences for ChIRP-Seq

Mrhl Oligo 1: 5’-AGTCAGATTACTGCTGGTCAGAACTAATAAACTCA-3’

Mrhl Oligo 2: 5’-CTGCTTCCTTCCTGGAATCAACAATAAAGCAGTTA-3’

Mrhl Oligo 3: 5’-ACTTCTTTCCAGTGACTGCAATTATCTTACAGAAGA-3’

Mrhl Oligo 4: 5’-TGAGTTTATTAGTTCTGACCAAGCAGTAATCTGACT-3’

Mrhl Oligo 5: 5’-TAACTGCTTTATTGTTGATTCCAGGAAGGAAGCAG-3’

Mrhl Oligo 6: 5’-TCTTCTGTAAGATAATTGCAGTCACTGGAAAGAAGT-3’

LacZ1: 5’-CCAGTGAATCCGTAATCATG-3’

LacZ2: 5’-TCACGACGTTGTAAAACGAC-3’

## Competing Interests

The authors declare that they have no competing interests.

## Acknowledgements

We thank Prof. Tapas Kumar Kundu (JNCASR, India) for providing E14TG2a mESCs and Prof. Maneesha Inamdar (JNCASR, India) for intellectual discussions. We thank Suma B.S of the Confocal Imaging Facility, Dr. R.G Prakash of the Animal Facility and Anitha G. of the Sequencing Facility at JNCASR, India. We thank Dhanur P. Iyer for initial help in standardization of RNA FISH technique. M.R.S Rao acknowledges Department of Science and Technology, Govt. of India for SERB Distinguished Fellowship and this work was financially supported by Department of Biotechnology, Govt. of India (Grant Numbers: BT/01/COE/07/09 and DBT/INF/22/SP27679/2018). Debosree Pal thanks University Grants Commission and JNCASR, India for her PhD fellowship.

