## Supplementary Figures and Tables for "LncRNA Mrhl orchestrates differentiation programs in mouse embryonic stem cells through chromatin mediated regulation"

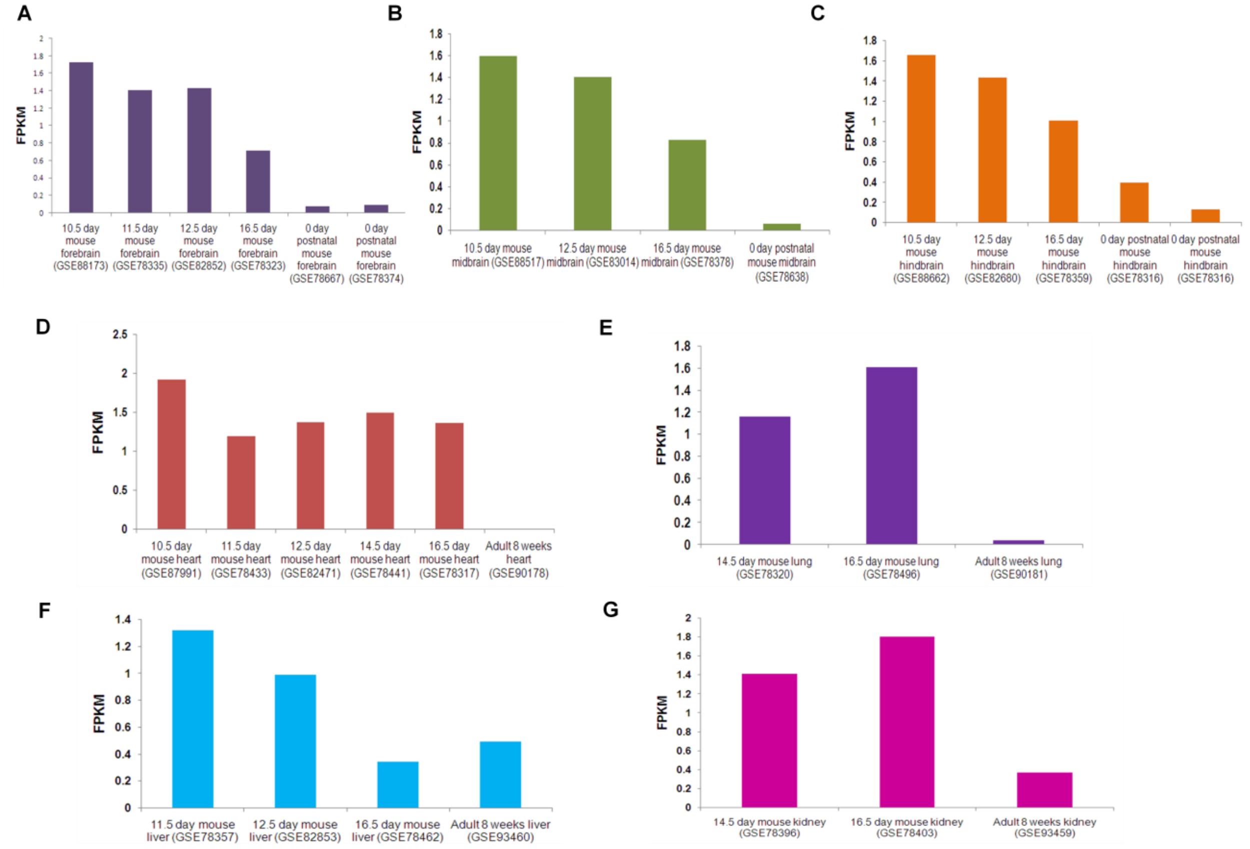


*Figure 1- supplementary figure 1: Analysis of Poly (A) RNA-Seq datasets from ENCODE database*

*Mrhl is expressed in various tissues at stages of E10.5-E16.5 and significantly down regulated in postnatal stages or adult stages. (A-C) Fore-, mid-, hindbrains; (D) heart; (E) lung; (F) liver; (G) kidney.*

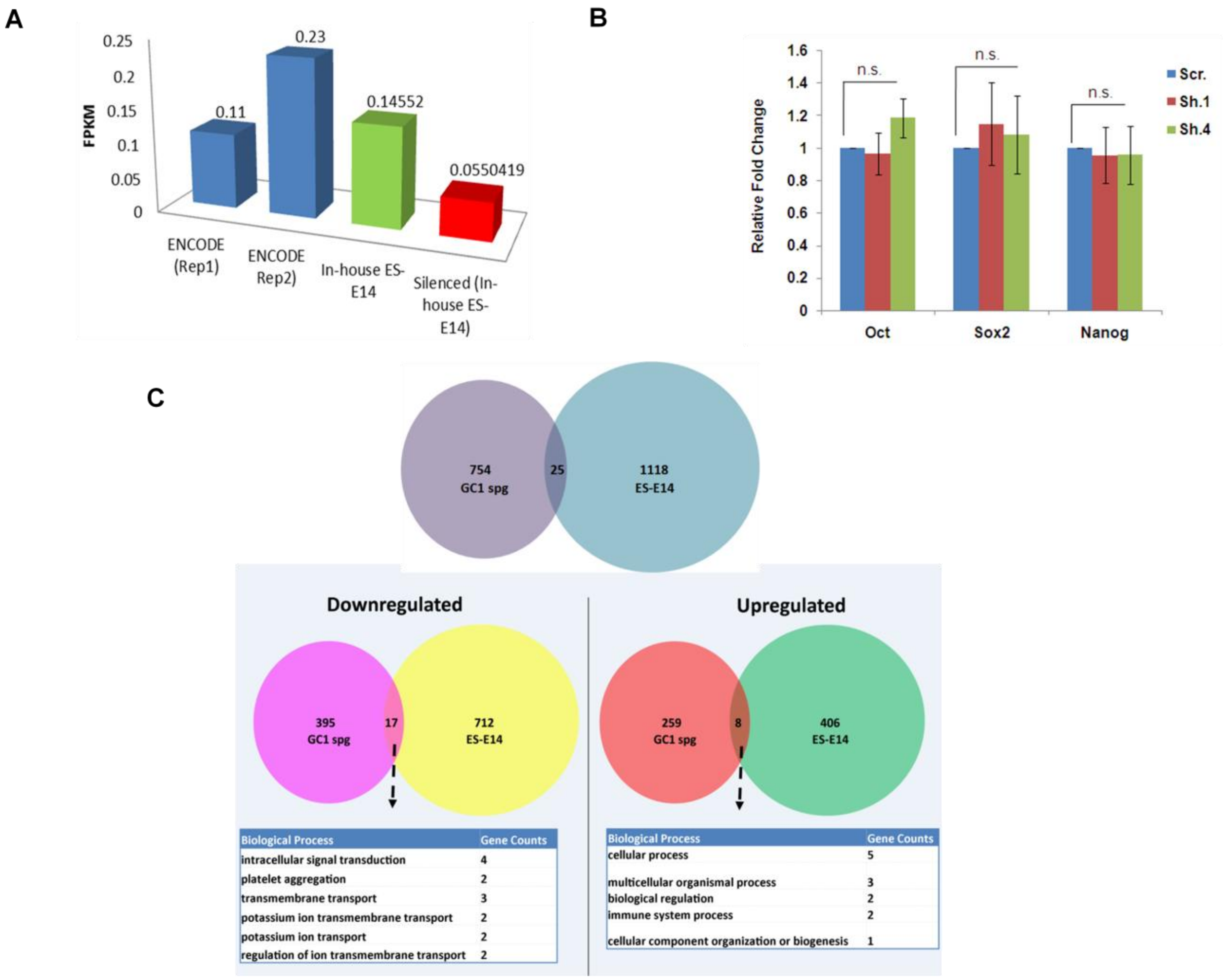


*Figure 2-supplementary figure 2: Analysis of DEG upon Mrhl depletion in mESCs. (A) Analysis of pluripotency markers in sh.1 and sh.4 treated mESCs as compared to scr.; (B) FPKM values for Mrhl in ENCODE versus in-house mESCs and scrambled versus sileneced mESCs; (C) Comaprison of DEG between mESCs and GC1-Spg spermatogonial progenitors. Error bars indicate standard deviation from three independent experiments. *p<0.05, **p<0.01, ***p<0.001, student's t-test.*

| GO-Slim Process | Obtained Number of Genes | Expected Number of Genes | P-Value | FDR | Fold Change | Obtained Over/under Expected |
|---|---|---|---|---|---|---|
| Neuromuscular synaptic transmission (GO:0007274) | 6 | 1.08 | 1.19E-03 | 3.22E-02 | 5.54 | + |
| Ectoderm development (GO:0007398) | 16 | 4.79 | 5.25E-05 | 4.27E-03 | 3.34 | + |
| Response to endogenous stimulus (GO:0009719) | 17 | 5.28 | 4.68E-05 | 5.72E-03 | 3.22 | + |
| Ion transport (GO:0006811) | 28 | 9.01 | 3.57E-07 | 8.71E-05 | 3.11 | + |
| cell-cell adhesion (GO:0016337) | 11 | 3.62 | 1.54E-03 | 3.41E-02 | 3.04 | + |
| Anion transport (GO:0006820) | 16 | 5.67 | 3.21E-04 | 1.30E-02 | 2.82 | + |
| Synaptic transmission (GO:0007268) | 20 | 9.03 | 1.25E-03 | 3.04E-02 | 2.21 | + |
| Cell-cell signaling (GO:0007267) | 29 | 13.55 | 2.54E-04 | 1.24E-02 | 2.14 | + |
| **Developmental process (GO:0032502)** | **60** | **34.89** | **6.44E-05** | **3.93E-03** | 1.72 | + |
| G-protein coupled receptor signaling pathway (GO:0007186) | 6 | 20.16 | 4.36E-04 | 1.52E-02 | 0.3 | - |
| immune response (GO:0006955) | 3 | 14.52 | 5.72E-04 | 1.75E-02 | 0.21 | - |

*Supplementary Table 1: List of GO from Fisher's exact test*

**A**

| Functions | Transcription Factors |
|---|---|
| Neuronal lineage | Atoh1, Tbr1, Dlx3, Lhx1, Vsx2 |
| Cardiac lineage | Myocd |
| Hematopoeitic and vascular lineage | Erg |
| Skeletal morphogenesis and osteoblast lineage | Runx2 |
| Myeloid and B-cell lymphoid lineage | Spi1 |
| Axis specification and positional identities | Hoxc4 |
| Mesoderm differentiation and limb patterning | Tbx2 |
| Trophoblast giant cells differentiation and cardiac morphogenesis | Hand1 |
| Epithelialization of somitic mesoderm and development of cardiac mesoderm | Mesp1 |
| Notch signaling | HeyL |
| Regulation of developmental processes | Tbx19 |
| Pancreatic development and maintenance | Pdx1 |
| Roles in epidermis and probable roles spermatogenesis, oocyte differentiation and embryo development | Bnc1 |

**B**

| Functions | Cell Adhesion and Receptor Activity related genes |
|---|---|
| Migration of epidermal cells and cerebellar development | Fat2 |
| Cell adhesion regulation during development | Col24a1, Col22a1, Lamc3 |
| Neuromuscular circuit development, axon guidance | Epha3 |
| Ca (+2) dependant cell adhesion and neurite growth in hippocampal neurons | Lrfn5 |
| Axon guidance and neuronal cell survival | Uncd5 |
| T-cell biology and neuron regeneration | Nav3 |
| NMDA receptor signaling and regulation of synapses | Dlg2 |
| Angiogenesis and spinal cord neurogenesis | Dll4 |
| Growth and differentiation of neuronal, glial, epithelial and other cell types | Nrg2 |
| Hematopoeisis | Flt3 |
| Acrosome expansion during spermatogenesis | Spaca1 |
| Muscle contraction and structural role in muscles | Mybpc3, Mybpc1 |
| Component of basement membranes; role in hearing and vision | Ush2a |
| Cellular proliferation and differentiation during retina and bone formation | Gdf6 |
| Osteogenesis and adipogenesis | Gdf10 |

*Supplementary Table 2: Functions of genes belonging to GO: 0032502. (A) Transcription factors; (B) Cell adhesion and receptor related genes (source: NCBI Gene and UniProtKB)*

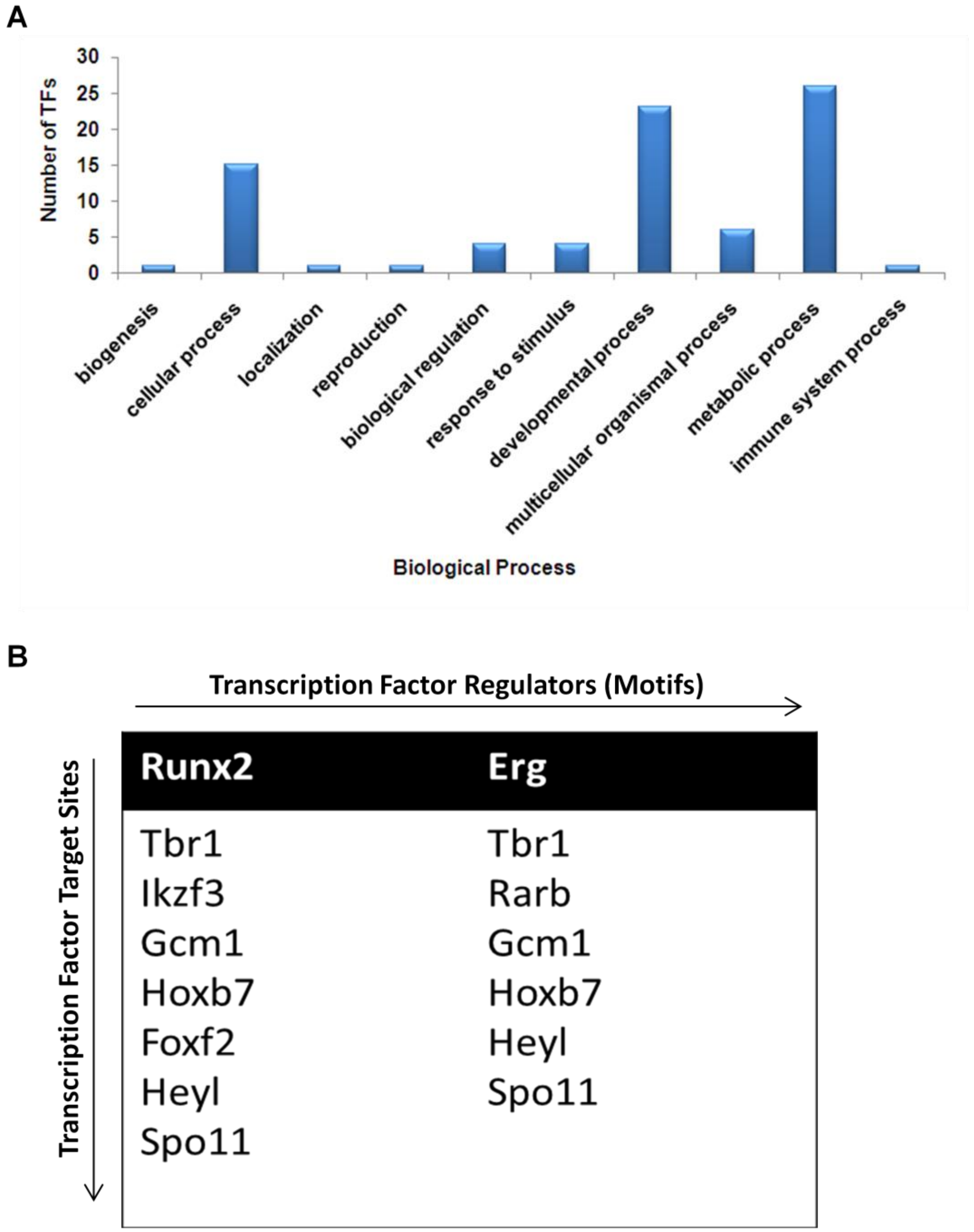


*Figure 3-supplementary figure 3: TF network analysis. (A) Gene ontology for perturbed TFs; (B) RUNX2 and ERG have the maximum number of binding sites on the target TFs.*

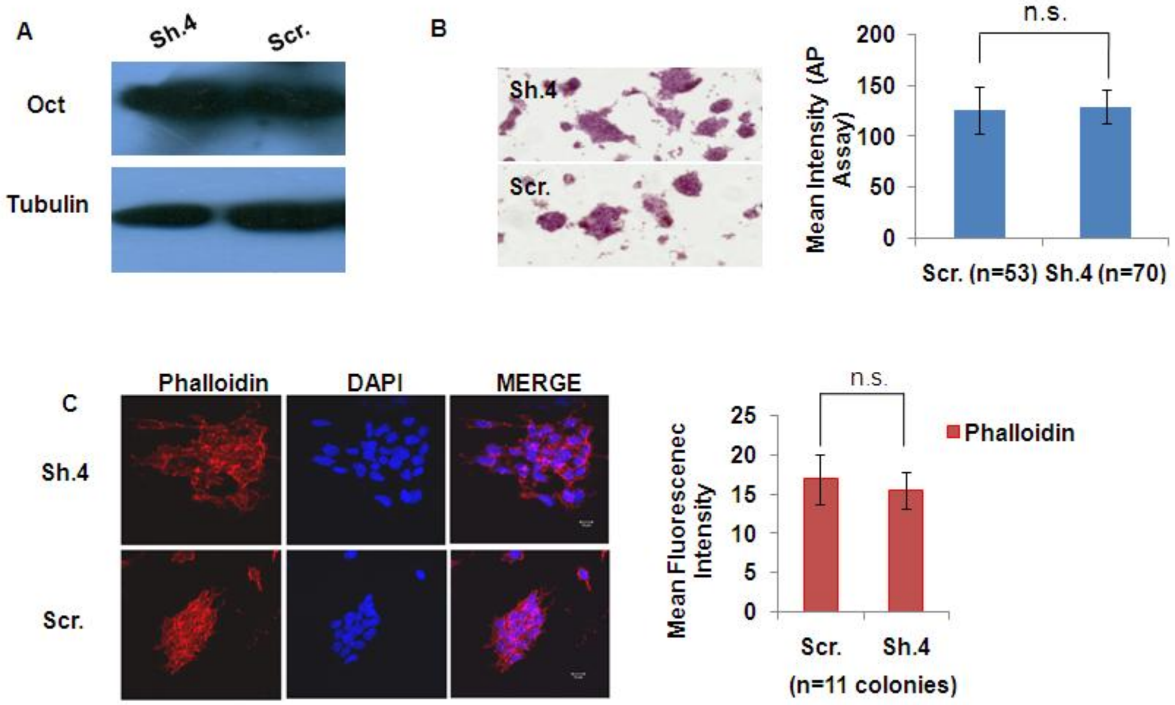


*Fig. 4-supplementary figure 4. (A) Oct4 western blot in stable knockdown versus scrambled control cells; (B) Alkaline phosphatase assay does not show any changes in pluripotency status of cells; (C) Phalloidin IF to validate cell adhesion status. Error bars indicate standard deviation from two independent experiments. *p<0.05, **p<0.01, ***p<0.001, student's t-test; Scale bar = 10 μm.*

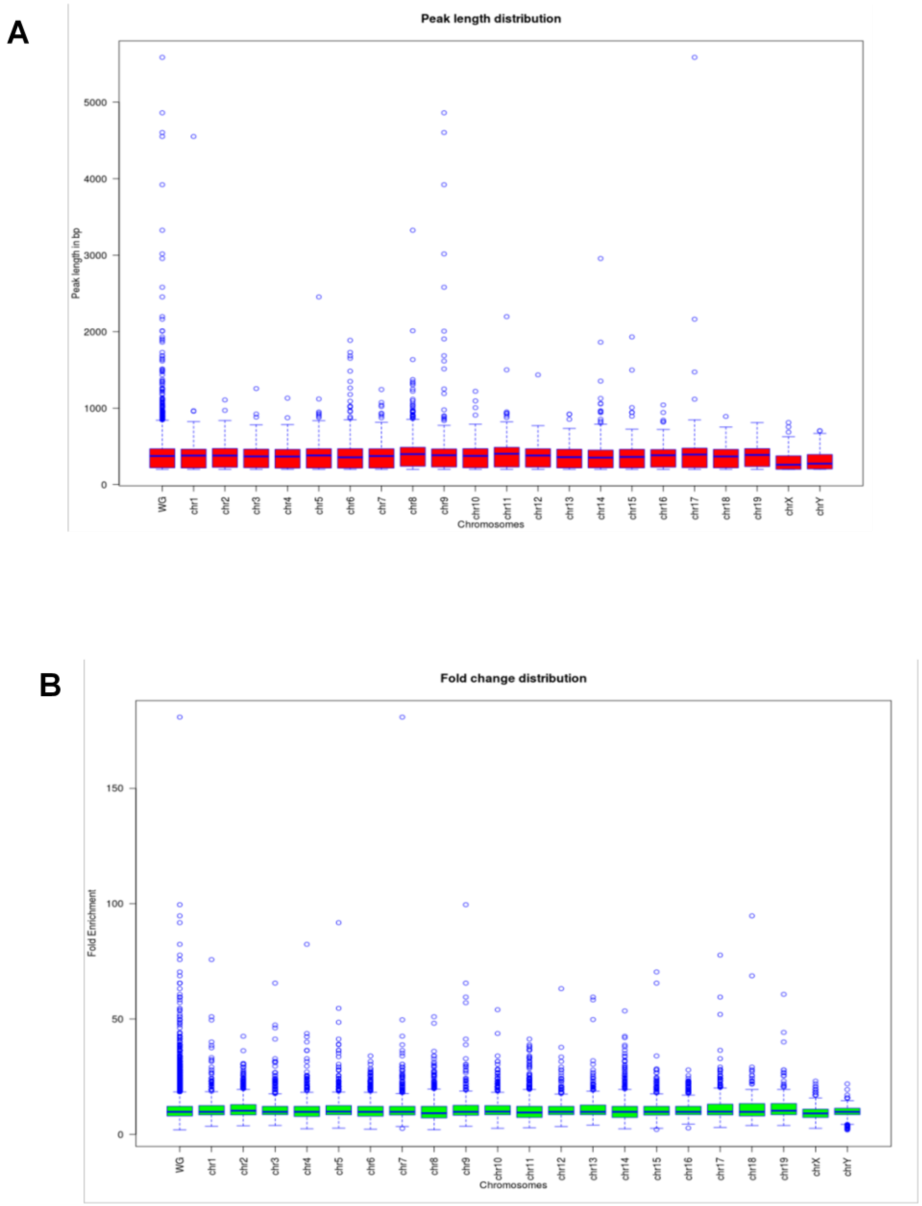


*Figure 5-supplementary figure 5: Distribution of ChIRP-Seq parameters across chromosomes*

*(A) Distribution of peaks lengths across chromosomes; (B) Distribution of fold change across chromosomes*

| ChIRP-Seq Shortlisted Target Genes | Functions |
|---|---|
| Six2 | Role in nephron progenitors during kidney development, craniofacial development and stomach organogenesis |
| Runx2 | Osteoblastic differentiation and skeletal morphogenesis |
| Dlx3 | Probable roles in the development of the ventral forebrain, craniofacial patterning and morphogensis |
| Hoxb7 | Determination of positional identities during development |
| Pou3f2 | Neuronal differentiation and reprogramming of somatic cells into neurons along with ASCL1 and MYT1L |
| Foxp2 | Implicated in the development of neural, gastrointestinal and cardiovascular tissues |

*Supplementary Table 3A: Development related TFs from overlap set of genes from RNA-Seq and ChIRP-Seq (source: NCBI Gene and UniProtKB)*

| Motif | Motif width | % of peaks | E-value | Number of occurrences | | | | | | Total % of peaks occupied by motifs |
|---|---|---|---|---|---|---|---|---|---|---|
| | | | | Exon | Intergenic | Intron | Non coding | Promoter | TTS | |
| Motif 1 | 15 | 21.46% | 6.2e-105 | 10 | 2701 | 1739 | 2 | 60 | 41 | 92.71 |
| Motif 2 | 15 | 28.16% | 4.9e-058 | 35 | 3504 | 2274 | 6 | 82 | 60 | |
| Motif 3 | 15 | 43.08% | 2.9e-021 | 80 | 5453 | 3350 | 10 | 134 | 83 | |

*Supplementary Table 3B: Occurrences of each of the 3 Mrhl motifs at the 21282 genomic loci*

| Motif ID | Alt ID | Sequence Name | Strand | Start | End | p-value | q-value | Matched Sequence | Potential Triplex Found |
|---|---|---|---|---|---|---|---|---|---|
| 3 | RVASAVAVDMASRMA | Pou3f2 | - | 95 | 109 | 0.000941 | 0.832 | GAAGAGGCAGACTCA | Yes |
| 1 | AGRAAGRRARRAARR | Foxp2 | - | 301 | 315 | 7.85E-10 | 1.28E-07 | AGGAAGGAAGGAAGG | Yes |
| 2 | RRRRRRMAGVMAGVV | Foxp2 | - | 300 | 314 | 1.74E-09 | 2.95E-07 | GGAAGGAAGGAAGGA | Yes |
| 3 | RVASAVAVDMASRMA | Foxp2 | - | 412 | 426 | 6.66E-09 | 8.28E-07 | AAAGAAAGAAAGAAA | Yes |
| 3 | RVASAVAVDMASRMA | RunX2 | + | 175 | 189 | 0.000179 | 0.0857 | TAATGAAAACAGGAA | Yes |
| 3 | RVASAVAVDMASRMA | Hoxb7 | + | 115 | 129 | 0.0000168 | 0.00898 | ACTGACAGAGAGACA | No |
| 3 | RVASAVAVDMASRMA | Hoxb7 | + | 174 | 188 | 0.0000929 | 0.0249 | AGAACAACAAAGACA | No |

*Supplementary Table 3C: Motif consensus, matched sequences and triplex formation prediction by Mrhl at target loci*
