## Supplementary File 6 for "LncRNA Mrhl orchestrates differentiation programs in mouse embryonic stem cells through chromatin mediated regulation"

**Motif (1/2/3)**

**Triplex Target Site (TTS)**

**>Pou3f2 (Motifs 3 is present)**

GAGCCCCCTTGACGCTCACCTCGATGGAGGTCCGCTTTTTCCGTTTGCGCCCTTGCGCTGCGATCTTGTCTATGCTGGTGGGGCTGCCCGAGGATGAGTCTGCCTCTTCCAACCACTTGTTCAACAAAGGCTTCAGCTTGCACATGTTCTTGAAGCTCAGCTGCAGGGCCTCAAACCTGCAGATGGTGGTCTGCGAGAACACGTTGCCGTACAGGGTGCCAAGCGCCAGCCCCACGTCTGCTTGAGTAAATCCGAGTTTGATCCGCCTCTGCTTGAATTGCTTGGCGAACTGCTCCAGGTCGTCTGAGGTCGGCGTGTCCTCGTCCGAGTGCGGGTCGTGGTGCGCGCCTGGGTGGCCCGGTGGGCCTTGTGGGGGAGGTGGCGGGGGCGGTTGCTGGTGTGGGTGAGAGTGCGGATGCGGGTGGTGGTCTGCATGGTGTGGCTCATCGTGGGCGT

(TGAGTCTGCCTCTTC Motif 3)

**>Foxp2 (Motifs 1, 2 and 3 are present)**

CTTCTTTTCTCTCTAGTTAAGAAACTCCTGGCCCCTAGTGACTCCTGAGATAACCATGCATTTCTCGGGCCACTCCTGATCACTCTCTTCCACTTGTCTGCTAGCCAGCAAGGCCTGCGGCTTCCCAATGGATCCGTCCTGGTTTTACTTTCCTTCCACTTTGGGAAGTGCTCCTTGTTCTGTGTGTGGGTGTCACAGCTTGCAGCCCCTGGGAGAGGGCATGTGGGAGGGGCACAGACACCTTGGTGAGAAAGGGTTTCTCCCTAGCCTTTTTTTTTCTTTCTCCTTCCTTCCTTCTCTCCT**TCCTTCCTTCCTTCC**TTCCTTCCTTCCTTCCTTTTTTCTTCCTTCCTTCTCTCCTTTCTTCCTTCCTTCCTTCCTCTCTTCCTTCCTTTTTTCTTATCTCTCTCTCTCTTTCTTTCTTTCTTTCTTTCTTTCTTTCTTTCTTTCTTTCTTTCTTTCTTTCAAACCCATGAGTCTTAAGTGGCTGGGATCAGTGCTGCGAGTGAGGGAAGATCAACTCCAGCCTCCACTACATGCTCTTGGAGAGGCATTTGTGACTGTAGGTGCTGGCTAGACTCTCCAGGTGATCACAGGCACTGTAGGTGCACCTCTCCCTACCCTCAGGCCAGAAGTGCCCTGAATCTCCATCTTAACGAGGGCCCGCACCCAACTAGTGCTTGGACAGGAAAAAA

CCTTCCTTCCTTCCT Motif1

**TCCTTCCTTCCTTCC** Motif2

TTTCTTTCTTTCTTT Motif3

**>RunX2 (Motif 3 is present)**

TGCCCCCACCCACCATCACAGTCATCCGTTCCATGCCACTCCTGGTTACCATCACACTAGGAAGAAATCTAACATGCAAATTCAGAGTGGCGTGGATAAATGGCAAAAAATGCCTAGGAAATTGGTCTGCTCGCCTTTATAATGTTTGTTGAAAAATCCTCCATCGCTCCCAACTAATGAAAACAGGAAGCTCTATTCATAAATGTGAAATTCACTGCCTATGATATATAATCATCCTAA

TAAGAAAATGAGCTCTAGACATAC

TAATGAAAACAGGAA Motif3

**>Hoxb7 (Motif 3 is present)**

CAGGCGAGGCCGCCAGACCTACAGTCGCTACCAGACCCTGGAGCTGGAGAAGGAGTTCCTATTTAATCCCTATCTGACTCGCAAGCGGAGGATCGAGGTATCGCACGCGCTGGGACTGACAGAGAGACAGGTCAAAATCTGGTTCCAGAATCGGAGAATGAAGTGGAAAAAGGAGAACAACAAAGACAAGTTTCCCAGCAGTAAATGCGAGCAGGAGGAGCTGGAGAAAGAGAAGCTGGAGCGGGCACCAGAGACCGCCGAGCAGGGCGATGCGCAGAAGGGTGACAAGAAG

ACTGACAGAGAGACA Motif 3

AGAACAACAAAGACA Motif 3

No triplex found within the Mrhl occupied region
